# PEPR-GNN: Perturbation-Enhancer-Promoter-RNA Graph Neural Networks for Multiome Perturb-Seq modeling of regulomes

**DOI:** 10.64898/2026.05.05.722311

**Authors:** Zachary Markham, Boxun Li, Landon Nguyen, Lei Wang, Nikhil V. Munshi, Gary C. Hon

## Abstract

Cellular reprogramming is a complex interplay between perturbations and regulatory elements, culminating in gene expression changes. Current computational approaches do not explicitly model these regulatory interactions. Here, we performed combinatorial reprogramming with cardiac transcription factors, followed by Multiome Perturb-Seq to measure perturbations, open chromatin, and gene expression in individual cells. We then developed PEPR-GNN (Perturbation-Enhancer-Promoter-RNA Graph Neural Network), a theoretical and computational framework to model regulome responses during complex genetic perturbations. By statistically associating gene regulatory relationships, PEPR-GNN organizes genes into regulomes with shared gene regulatory responses to reprogramming, including easy-to-reprogram cardiac genes, difficult-to-reprogram fibroblast genes, and context-specific genes where the impact of a reprogramming factor depends on the presence of others. Finally, we use PEPR-GNN for *in silico* modeling of how genetic modifications of enhancers can be used to tune gene responses to reprogramming. Overall, through the use of causal perturbation information and an enhancer-aware regulome model of gene regulation, PEPR-GNN can effectively model complex cellular responses to perturbation.

**Highlights:**

- Multiome Perturb-Seq of GHMT reprogramming in MEFs with RNA/ATAC-Seq readout.
- PEPR-GNN: a computational framework to model perturbation-induced regulomes.
- PEPR-GNN aids the interpretation of regulomes by diverse reprogramming responses.
- PEPR-GNN enables in silico perturbation to tune gene responses to reprogramming.

## Introduction

Ectopic induction of transcription factors (TFs) is sufficient to reprogram the epigenetic, transcriptional, morphological, and function state of cells^1–4^. One approach to convert fibroblasts to cardiomyocyte-like cells is by induction of GHMT (Gata4, Hand2, Mef2c, Tbx5)^5^. Interestingly, induction of GHMT with additional TFs, even those that directly trigger cardiac gene expression in other contexts, does not always improve reprogramming efficiency. This strategy can even yield greatly decreased reprogramming efficiency^6^. Notably, some cardiac genes are more susceptible to these unexpected combinatoric effects than others^7^. These difficulties indicate that, in the context of cardiac reprogramming, there is a complex interplay between ectopically expressed TFs and the cell’s context that may be antagonistic to complete reprogramming^6,8,9^. Understanding this interplay will be important to developing improved reprogramming cocktails and to understand the mechanisms of cellular reprogramming.

Here, we address this problem experimentally, conceptually, and computationally. Experimentally, while reprogramming is the product of how TFs (perturbations) impact the cell’s epigenome (mechanism) and transcriptome (phenotype), previous functional genomic studies have only analyzed these 3 components systematically in isolation or in pairs. Thus, it remains difficult to distinguish correlative and causative associations relevant to reprogramming. In this study, we resolve this issue by systematically measuring all 3 components using Multiome Perturb-Seq of GHMT reprogramming cells. Conceptually, this approach enables a framework to model each gene’s regulome as a flexible graph of the 3 components acting in concert to determine the gene’s behavior. Computationally, since multiple regulatory mechanisms can be acting on a gene at once, it is essential that any model of gene response to perturbation can handle overlapping and non-linear relationships between the gene’s parts. Thus, we take advantage of developments in Graph Neural Networks (GNN) to model gene behavior during reprogramming. PEPR-GNN combines regulome graph construction with GNN modeling to focus on the regulatory relationships within each gene, giving not only regulatory-element-specific predictive power but also insights into how genes reprogram.

## Results

### A regulome model for reprogramming with Multiome Perturb-Seq

Conceptually, we model transcriptional reprogramming as a cascade from induction of exogenous TFs to the binding and activation of enhancers, leading to the activation of promoters and changes in gene expression. Each of these steps is carried out by distinct cellular mechanisms. For a given gene, it is the culmination of these components that gives rise to the variety of gene regulatory responses seen in different cell-contexts or with different perturbations. Thus, to understand and model how a gene will respond to perturbation during GHMT reprogramming, we reason that it is necessary to model how a gene’s regulatory components relate to each other (Figure 1A).

**Figure 1:**
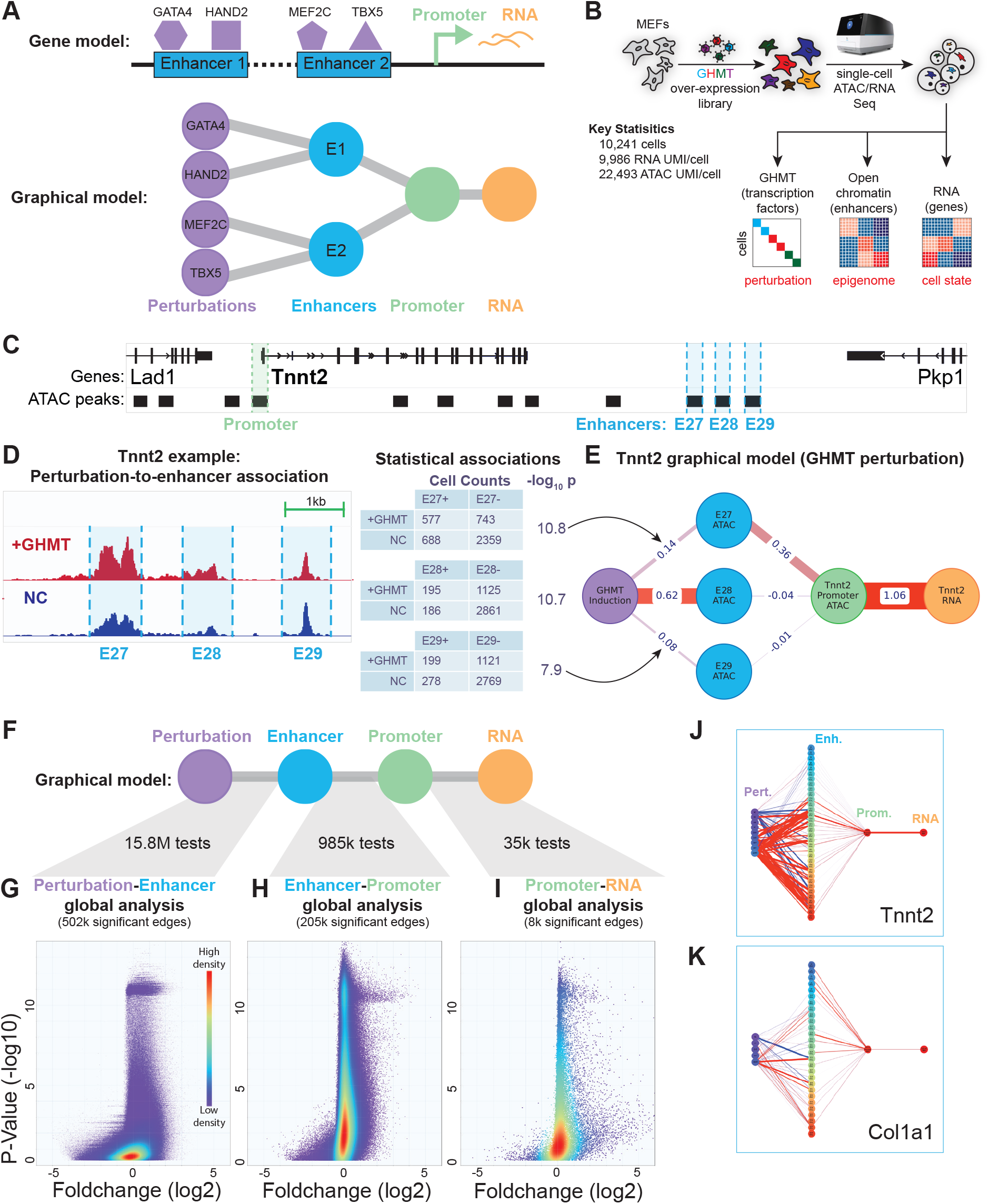
A genome-wide regulome model for reprogramming with Multiome Perturb-Seq. A. (top) Regulome model illustrating the action of GHMT factors binding enhancers to induce changes in promoter accessibility and gene expression. (bottom) Graphical model represention. B. Mouse embryonic fibroblasts (MEFs) were transduced with a lentiviral library for combinatorial induction of GHMT. Multiome Perturb-Seq readouts for each cell include perturbation, open chromatin, and expression information. C. Genome browser snapshot near the Tnnt2 gene, with regulatory elements labeled. D. Example perturbation-enhancer associations for the Tnnt2 regulome: (left) Open chromatin profiles for cells with detection of all 4 GHMT factors or no factors; (right) 2×2 contingency tables illustrating the calculation of associations. E. A subset of the Tnnt2 regulome after statistical associations, focusing on one perturbation (GHMT) and three enhancers. F. Regulome schematic for systematic testing. G. Genome-wide statistical association analysis for perturbation-enhancer edges. Density of points is indicated. H. Genome-wide statistical association analysis for enhancer-promoter edges. I. Genome-wide statistical association analysis for promoter-RNA edges. J. Example regulome architectures for cardiac gene Tnnt2. K. Example regulome architectures for fibroblast gene Col1a1.

To address this question, we performed GHMT reprogramming on mouse embryonic fibroblasts (MEFs), followed by a single-cell multiomic readout (Figure 1B). For each cell, we measure three modalities: the expression of exogenous GHMT transcripts, open chromatin status (ATAC-Seq), and expression status (RNA-Seq). We performed this experiment at moderate MOI to ensure the detection of cells with each distinct combination of reprogramming factors (Table S1).

For a given gene, our regulome model considers four components of its regulation: 1) The perturbation state of the cell, determined by which exogenous GHMT transcripts are detected; 2) Enhancer open chromatin status and 3) promoter open chromatin status, determined by the abundance of ATAC-Seq reads at these regulatory regions; and 4) The transcriptional state of the cell, determined by RNA-Seq expression.

We illustrate the example of Tnnt2, a cardiac gene induced by GHMT reprogramming^10^. In cells receiving exogenous copies of all 4 GHMT factors, we observed open chromatin peaks at the Tnnt2 promoter as well as nearby regulatory elements dubbed E27, E28, and E29, approximately 30-kb from the Tnnt2 promoter (Figure 1C). To quantify the relationship between GHMT induction and these three elements, we partitioned cells between those with all 4 GHMT factors induced versus negative control (NC) cells, which had no detection of GHMT factors. Notably, we found that cells with GHMT perturbation had 54% more enrichment for open chromatin than negative control cells at E28 (p<1e-10, log(E28_fc_) = 0.62). E27 and E29 had significant but weaker enrichment (log(E27_fc_) = 0.14, p<1e-10; log(E29_fc_) = 0.08, p<1e-7) (Figure 1D). The relationship between any pair of components (perturbation, enhancer, promoter, RNA) can be quantified similarly (Figure S1A, also see processed data in GEO: GSE329363). Combining these results gives a multi-step network connecting perturbations to regulatory elements and ultimately to Tnnt2 expression (Figure 1E). By examining each relationship type in context with the others, it becomes apparent that E27 is the most plausible element by which GHMT induces Tnnt2 expression. In contrast, while E28 gains more open chromatin upon GHMT induction, its association with TNNT2 expression is significantly lower. These observations motivate the extension of this regulome model to gain predictive insights into reprogramming.

### Genome-wide modeling of regulomes

Generalizing the example above, we model each gene as a graph, containing 4 types of nodes representing each of the components of our regulome model: Perturbation nodes (i.e., G, GH, GHMT), Enhancer nodes (i.e., E27), Promoter nodes (i.e., the Tnnt2 promoter), and RNA nodes (i.e., Tnnt2 RNA). Edges represent statistical associations between nodes. For example, the edge connecting E27 with the Tnnt2 Promoter is an “enhancer-promoter” type and has an associated fold-change and p-value. To construct regulome models globally, we extended the Tnnt2 example to all genes. In Figure 1F, we illustrate three relationships, or edge types, that link these components together.

- <u>Perturbation-enhancer edges.</u> Genome-wide, we consider linkages between all 16 perturbation states in our dataset to all detected enhancers (∼15.8 million edges) (Figure 1G). We found >500,000 significant edges, mostly enhancers that gain accessibility upon TF induction. The strongest edges connect reprogramming cocktails with 3 or 4 TFs, which have a more pronounced effect on enhancer activity across the genome than milder 1-2 TF perturbations (Figure S1B). Edges with negative fold change often connect fibroblast enhancers that decrease activity upon perturbation.
- <u>Enhancer-promoter edges:</u> Our analysis of nearly 1 million possible enhancer-promoter edges yields >200,000 statistically significant associations (Figure 1H). Consistent with the role of enhancers to activate gene expression, most significant edges have a positive fold-change.
- <u>Promoter-RNA edges:</u> We consider a total of 35,038 promoter-RNA edges, of which just over 8,000 are statistically significant (Figure 1I). This high proportion of significant edges (∼23%) is due to the strong link between promoter activity and gene expression. As we model a separate graph for each unique promoter, genes with alternative promoter usage can have unique structures. (Figure S1E).

The set of regulome graphs we constructed represent diverse structures. We illustrate several examples for canonical cardiac genes (Tnnt2, Hcn4, and Ryr2) and fibroblast genes (Mfap4, Col1a1, and Ccdc80), which undergo active regulatory reprogramming (Figure 1J-K, S1C-D). We observe that the node representing induction of all 4 GHMT factors is the most connected perturbation node, consistent with the synergistic activity of this combination to activate enhancers and induce cardiac genes^5,10^. In contrast, regulomes for fibroblast genes have distinct structures consistent with repression. For example, strongly promoter-linked enhancers, such as E13 of Mfap4, are negatively associated with GHMT. Also, while some fibroblast gene regulomes have enhancers positively associated with GHMT, these enhancers are often not significantly linked to promoter activity, suggesting that they do not maintain the expression of these fibroblast genes.

### Functional embedding of regulomes with graph neural networks

To systematically identify regulatory patterns across regulome graphs genome-wide, we developed a flexible and robust graphical neural network (GNN) architecture^11^. One key consideration in our design was the use of heterogeneous GNNs to model the different node and edge types in our regulome graphs, due to the differing biological significance attributed to each. This approach also allowed us to incorporate different biological features into each node type, which could aid modeling (Figure 2A). Perturbation nodes were represented as a binary string encoding the 16 combinations of 4 GHMT factors. Enhancer and promoter nodes also include genomic sequence information, encoded as the frequency of 6-mer motifs in the genomic region, a length which we found to strike a balance between specificity and redundancy^12^. This encoding gives a simple but information-rich representation of each node’s potential TF binders. Lastly, RNA nodes encapsulate changes in expression for each perturbation.

**Figure 2:**
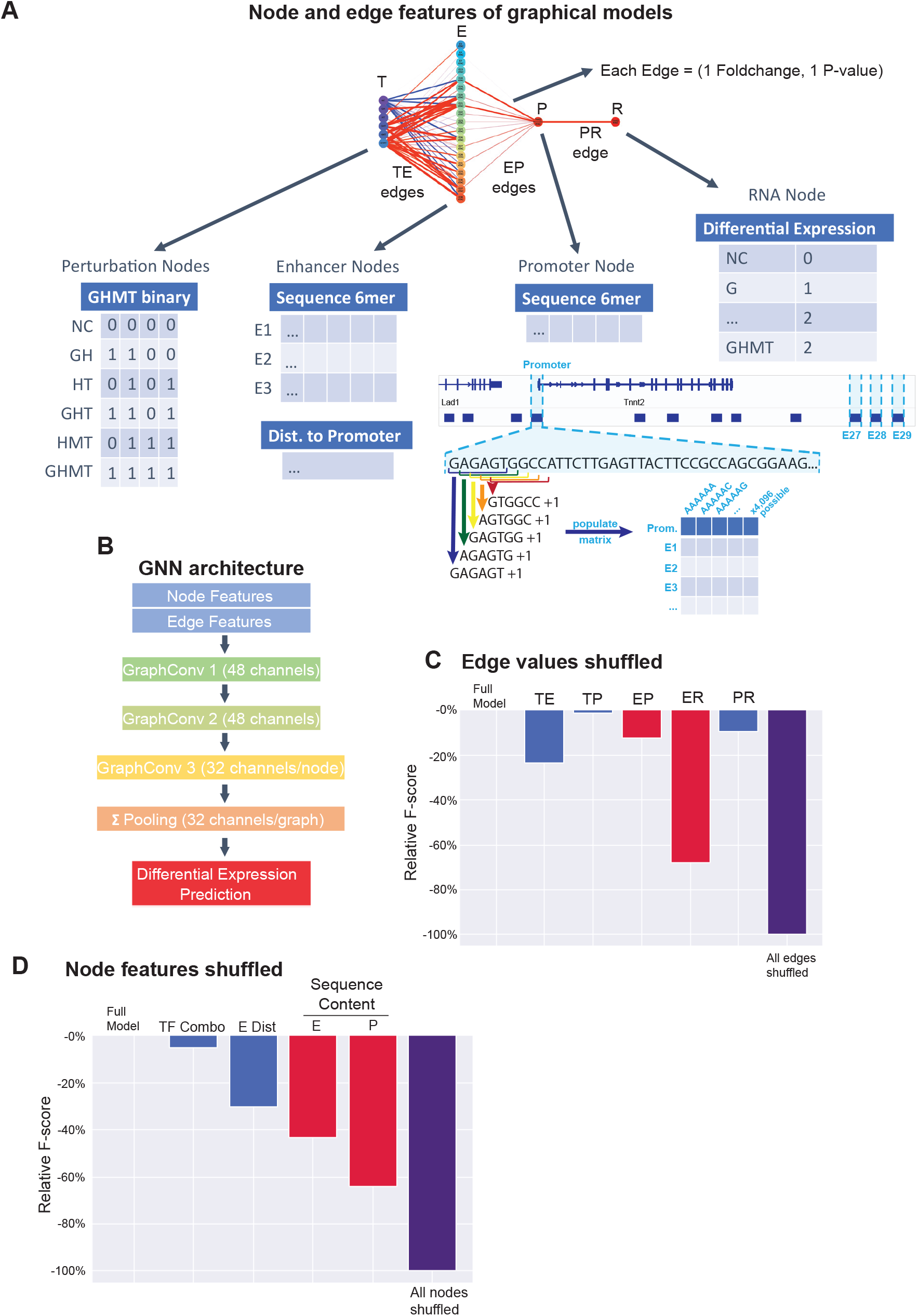
Structuring Gene Graphs as inputs for a GNN Architecture. A. Structure of graphical models input into GNN architectures. Perturbation nodes are represented as a binary string encoding TF combinations. Enhancer/promoter nodes contain sequence information encoded as the 6-mer count matrix tiling the corresponding open chromatin peak, independent of motif placement or local sequence context. Enhancer nodes also contain information on distance from the promoter. RNA nodes contain a trinary string indicating if they are positively, neutrally, or negatively, associated with each of the 16 combinatorial TF perturbations. Edges contain statistical information including p-values and fold changes of association between two nodes. B. Input graphs are sent through a GNN architecture consisting of 3 GraphConv layers with a final summation-pooling layer to create a single graph-level output vector. Normalization is performed at each layer. The processed output vector contains 16 values, corresponding to the predicted responses to each of the 16 perturbations. C and D. All edge (C) and node (D) features are randomly shuffled, one type at a time, to assess their impact on model performance. Performance is measured in terms of F-score, which is determined by prediction accuracy and precision. All values are scaled relative to the performance of shuffling all edge or node features respectively.

The unique properties of our dataset, with heterogeneous node types and multiple weighted edges per node-pair, made existing methods inappropriate. Thus, we developed a new approach using the GraphConv algorithm from Pytorch-Geometric^13^. After an extensive parameter search, our final architecture consisted of three layers, with Dropout^14^ and ReLU^15^ performed after each convolution (Figure 2B). A BCE loss-function^16^ was used for training the network to predict a graph’s transcriptional response to each possible combination of GHMT factors. We reasoned that a gene’s response to a given perturbation is an emergent property of the regulatory elements that constitute it and how these elements interact. Therefore, a network tasked with predicting a gene’s response may be able to learn the direct or derived-features of the gene graph that underlie this property.

After training and optimization, the Perturbation-Enhancer-Promoter-RNA GNN (PEPR-GNN) model could generate predictions for how any gene responds to any GHMT perturbation. We first used the trained model to perform feature importance analysis by randomizing the values of each feature type, and observing the effect on performance. The most informative features of the model are the edges connecting from enhancers (Figure 2C) and sequences at enhancers/promoters (Figure 2D).

The trained PEPR-GNN model can be used to convert diversely structured regulome graphs to uniformly-sized vector representations. After dimensionality reduction (Figure 3A), the resulting 2D tSNE visualization represents regulome responses across all genes during GHMT reprogramming. Coloring graphs according to how expression levels change during GHMT induction (Figure 3B) illustrates co-clustering of cardiac genes like Tnnt2 (right side), which are up-regulated during cardiac reprogramming. This co-clustering indicates that easily-reprogrammed genes share architectural similarity in their regulome graphs. Likewise, fibroblast genes like Fap are down-regulated during cardiac reprogramming and also co-cluster (bottom left). Finally, other genes that are not specific to cardiac or fibroblast states, such as Top2a, are found closer to the middle region.

**Figure 3:**
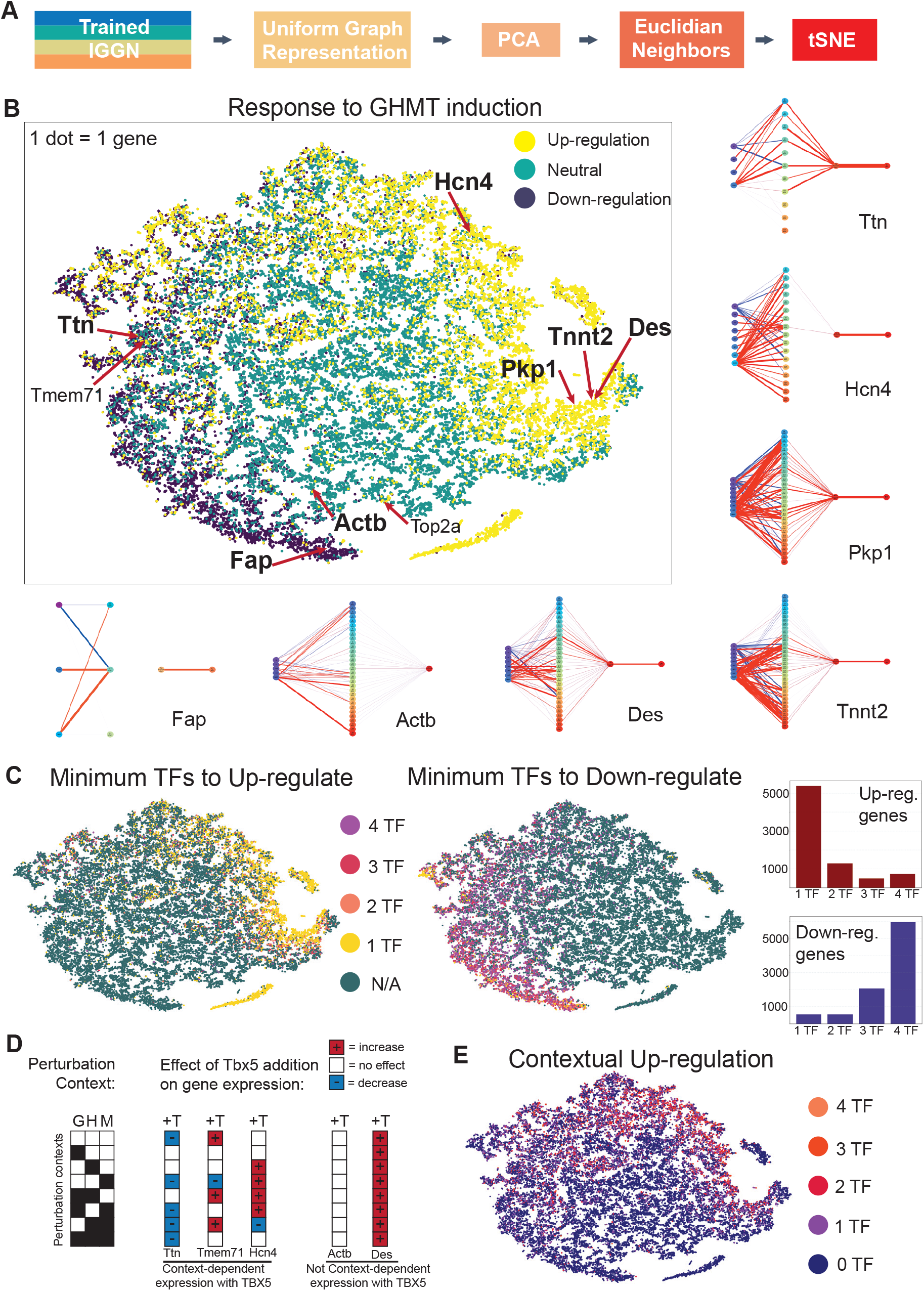
Functional embedding of regulomes with graph neural networks. A. Pooled model outputs can be used as features for tSNE embedding, placing every graph on a single manifold organized by features important to the model. B. (top left) Shown is a 2-dimensional tSNE embedding of gene graphs, colored according to GHMT response. The locations of notable genes are indicated. (right and bottom) Regulome models corresponding to key cardiac and fibroblast genes. C. Gene graphs are colored according to the minimum number of GHMT factors needed to up-regulate the gene. The cardiac genes on the right of the embedding can be robustly induced by even a single GHMT factor, and are “easier” to reprogram. D. Gene graphs are colored according to the minimum number of GHMT factors needed to down-regulate the gene. The fibroblast genes on the left of the embedding require the induction of many GHMT factors to repress gene expression, and are “harder” to reprogram. E. An example where Tbx5 exhibits contextual regulation of Ttn, Tmem71, and Hcn4 depending on the context of GHM factors expressed in a cell. In contrast, Tbx5 does not exhibit contextual regulation of Actb and Des. F. Gene graphs are colored according to the number of TFs that exhibit contextual up-regulation. The graphs on the top of the embedding exhibit the most contextual regulation.

### Interpreting regulomes by diverse reprogramming responses

We used the regulome embedding to gain insights into how gene regulatory interactions are restructured during GHMT reprogramming. First, we sought to determine the impact of overlapping enhancers, as there are many examples of the same enhancer node being present in multiple distinct gene graphs. For example, Tnnt2 and Pkp1 share 25 enhancer nodes, giving the two graphs a ∼60% overlap. Despite this degree of enhancer overlap, the nearest neighbor of Tnnt2 in the embedding is Des, with Pkp1 being the 173rd nearest neighbor (Figure 3B). Consistently, Des expression more closely matches Tnnt2 expression during reprogramming: Des is expressed in ∼77% of Tnnt2-positive cells while Pkp1 is expressed in only ∼6% of Tnnt2-positive cells. This suggests that PEPR-GNN is deriving features about each graph by considering the broader context of a graph’s regulome instead of just considering which elements are present.

Given that the embedding successfully separates graphs according to their response to GHMT induction, we next wanted to see if regulatory responses to intermediate perturbations (subsets of GHMT) were also spatially resolved. We colored the graphs by the minimum number of TFs needed to induce expression (Figure 3C, Figure S3A). For example, since Gata4 alone is sufficient to up-regulate Tnnt2, we assign Tnnt2 a value of 1. In contrast, since no single TF is sufficient to activate Hcn4 but the combination of Hand2 and Mef2c can, we assign Hcn4 a value of 2. We observe a cluster of genes near Tnnt2 that can be up-regulated by even a single TF (right side). Adjacent to this cluster is a zone of genes that require 2 or more TFs to be induced. Mirroring this observation, we observe a cluster of fibroblast genes near Fap that can be down-regulated by GHMT. However, one interesting asymmetry between up-regulated and down-regulated genes is that while 1-2 factors are often sufficient to trigger up-regulation, the vast majority of down-regulated genes require a minimum of 3-4 TFs (Figure 3C). This is consistent with previous studies and shows that cardiac and fibroblast gene pathways need not reprogram in lock-step with each other^17,18^. This observation also supports the idea that repression of the starting cell identity is a greater barrier to reprogramming than turning on target cell fate.

In contrast to the relatively simple regulation above, we observe cases where the impact of a given TF on a gene’s expression depends on the presence of other TFs induced in the cell (which we refer to as the cell’s context). For example, addition of TBX5 in the cell context of HAND2 alone or MEF2C alone induces the expression of Hcn4, while addition of TBX5 in cells with both HAND2 and MEF2C represses the expression of Hcn4 (Figure 3D). Thus, TBX5 contextually regulates Hcn4 (as well as other genes including Ttn and Tmem71), which contrasts with the uniform response that Tbx5 has on the expression of Tnnt2 and Fap across all GHM perturbation contexts (Figure 3C). Extending this analysis across all reprogramming TFs and all genes, we identify many instances of contextual regulation (for a comprehensive list, see our processed data in GEO: GSE329363). Notably, we observed that contextual up-regulation was disproportionately found in the top half of the embedding (Figure 3E). Our analysis generalizes previous studies showing that some genes are sensitive to the exact combination of reprogramming factors present in a cell^8^. Together, these observations underscore the difficulty of fully reprogramming the expression status of all genes in the genome to match a target.

### Synthetic perturbations of enhancer motifs to tune gene response

The trained PEPR-GNN model can be used to explore the effect of genetic alterations on gene expression *in silico*. First, we performed a synthetic screen: for a given gene and 6-mer motif pair, we made a copy of the graph and set the motif’s count for all the gene’s enhancers to 0 (Figure 4A). This allowed us to generate expression predictions spanning all possible gene-motif pairs, which were then scored by how greatly the predicted expression diverged from that of the wildtype gene. We observed that some genes were much more susceptible to expression alteration than others (Figure 4B). Ttn, Csrp3, and Myom2, were three cardiac genes scoring in the top 1% upon removal of the canonical binding motifs^19,20^ for the GHMT factors. This suggests that Ttn, Csrp3, and Myom2, are more sensitive to changes in GHMT activity at their loci. Importantly, the association between these genes and the GHMT motifs was not explicitly given to the model, and was instead learned during training. By contrast, Tnnt2 is among the lowest-scoring genes, and thus predicted to be less impacted by GHMT motif disruption. This is consistent with our observations that Tnnt2 can be robustly and redundantly induced by any GHMT factor. The presence of redundant mechanisms for Tnnt2 induction thus minimizes the impact of any single TF (or its motif’s disruption). The redundantly regulated fibroblast gene Fap behaves similarly.

**Figure 4:**
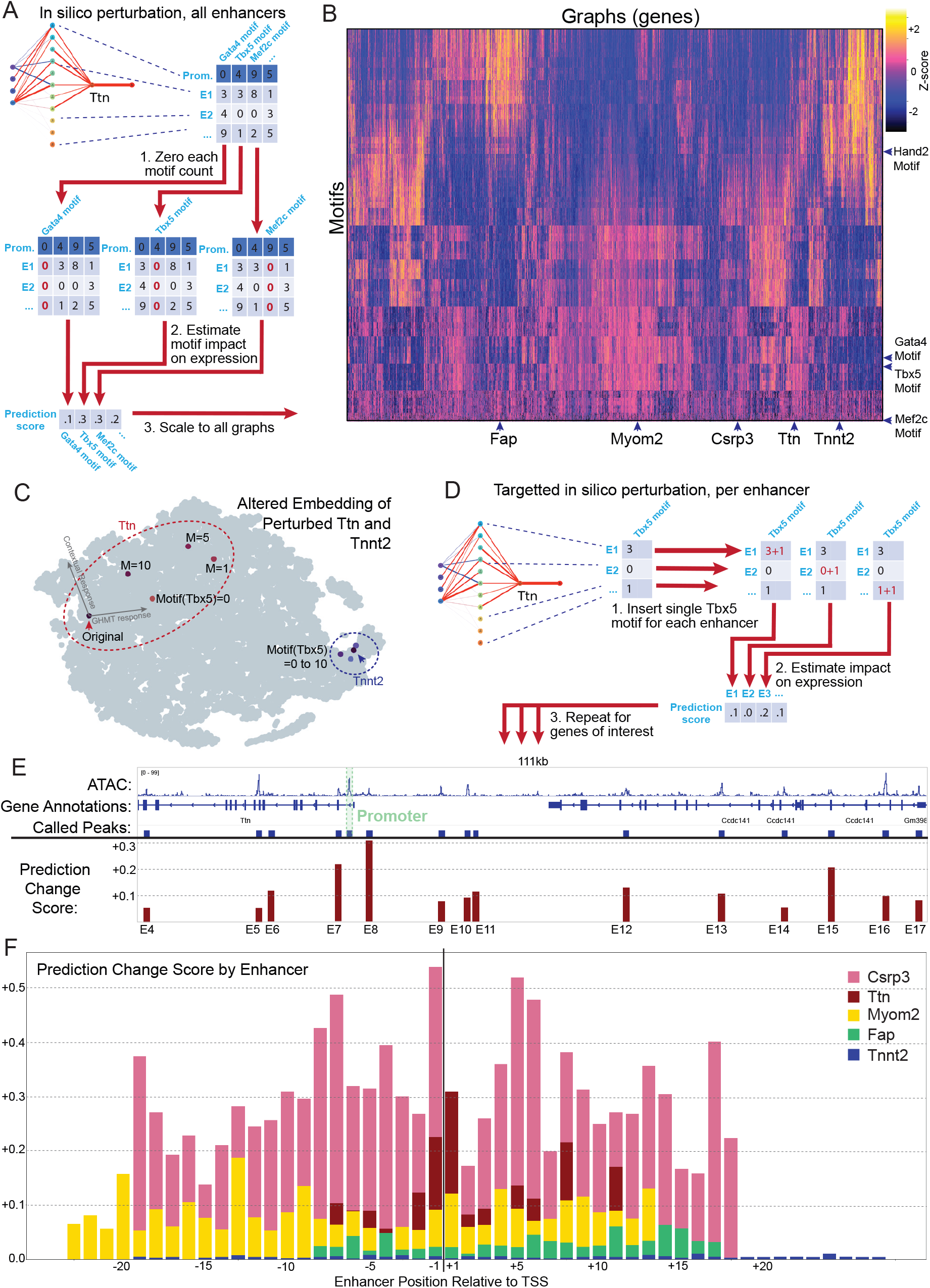
Interpreting regulomes by diverse reprogramming responses. A. Schematic of strategy to perform in silico screens to test the impact of enhancer motifs on gene expression using regulome models. (top) Regulome graph of Ttn with hypothetical motif counts for enhancers. We iterate through each 6-mer motif and set its motif count across all enhancers to 0. We then use the trained model to estimate the impact of this modified graph relative to the unmodified graph. We repeat for all graphs. B. The heatmap summarizes the impact of setting enhancer motif counts to 0 for all gene-motif pairs. Key reprogramming genes and motifs are indicated. C. For the Ttn regulome, we performed in silico perturbations to alter the Tbx5 motif count at all Ttn enhancers. We embedded the synthetic graphs to illustrate predicted impact. In comparison, we also repeated this process for Tnnt2. D. For each Ttn enhancer, we performed in silico mutagenesis to add one Tbx5 motif count and displayed the predicted change in expression. E. Genome browser snapshot of the Ttn locus with proximal enhancers and in silico predictions of Ttn expression change after addition of one Tbx5 motif to each enhancer. F. We repeated the in silico perturbation in (D-E) for Csrp3, Myom2, Fap, and Tnnt2. Shown are the predicted changes in expression from adding one Tbx5 motif count for each proximal enhancer.

To test the robustness of this idea, we performed extended *in silico* perturbation experiments for the Ttn-Tbx5 pair by varying Tbx5 motif counts at enhancers from 0 to 10. Across this range of values, the embedding location of Ttn shifts dramatically (Figure 4C). As embedding location is associated with a gene’s response to GHMT (Figure 3B, Figure S3A), this observation suggests that Ttn’s response to GHMT is substantially changed by the addition of Tbx5 binding sites. In contrast, a parallel analysis for the Tnnt2 regulome resulted in comparatively smaller changes in embedding location.

To increase resolution of *in silico* perturbation, we next performed a targeted synthetic screen on each enhancer in the Ttn regulome. We added a single Tbx5 motif to each enhancer node and scored enhancers by the resulting change in expression prediction (Figure 4D). The resulting score distribution is shown for enhancers around the Ttn locus (Figure 4E). The enhancer most impacted by addition of a single Tbx5 motif was E8, an open chromatin region ∼2 kb upstream of the Ttn promoter. Two other sensitive regions include E7 (∼1 kb downstream) and E15 (∼67 kb upstream). The remaining enhancers score much lower, suggesting only a small group of key enhancers account for Ttn’s overall sensitivity to motif perturbation. A similar analysis on Csrp3 and Myom2 showed high sensitivity enhancers (Figure 4F). Tnnt2 and Fap, by contrast, are much less responsive to perturbations of the Tbx5 motif at enhancers.

## Discussion

Our multiomic strategy to simultaneously detect changes in perturbagens, chromatin accessibility, and gene expression in individual cells is a general approach to learn about the interplay between gene regulation and cell state. Perturbations represent the axis of causality, while changes in open chromatin represent the regulatory mechanisms that culminate in the cellular phenotype of transcriptional change. What makes PEPR-GNN unique is its ability to model the statistical associations between each of these components (perturbations, enhancers, and genes). By applying PEPR-GNN to GHMT-mediated reprogramming, we garnered several insights. Consistent with previous studies, PEPR-GNN identifies “easy” cardiac genes (like Tnnt2 and Des) that can be robustly induced by any one of the GHMT factors. PEPR-GNN also identifies “difficult” fibroblast genes (like Fap) that require multiple GHMT factors for repression. Interestingly, we also identified a set of “contextual” genes (like Ttn and Hcn4) whose induction is highly dependent on the specific combination of GHMT factors induced in a cell. These results indicate that there is no single combination of GHMT factors that can successfully reprogram the expression of all cardiac genes, and imply that the choice of specific reprogramming cocktails should be application-specific and depend on the context-specific genes that need to be induced. It also suggests that adjusting the cell context of fibroblasts (established by its endogenously expressed TFs) could improve reprogramming. Alternatively, our in silico screens with PEPR-GNN show that genetically modifying specific enhancers to add or remove motifs could provide new capabilities to tune the expression of targeted reprogramming genes.

One key feature of PEPR-GNN is its incorporation of enhancer information to model reprogramming. Consistent with previous publications showing dynamic enhancer landscapes during reprogramming ^10^, we observe 41,571 distinct candidate enhancers statistically associated with at least one GHMT combination, accounting for more than one fourth of observed ATAC peaks across the genome. We even observe widespread enhancer activation near fibroblast genes, which are, nonetheless, typically repressed during cardiac reprogramming. This seeming contradiction can be resolved by asking what newly active enhancers do in these different contexts. PEPR-GNN is able to resolve that only 7,860 enhancers (less than one fifth of those activated) are associated with altered promoter chromatin status or gene expression. Thus, while GHMT factors flood the genome with chromatin changes, only a fraction go on to alter gene expression. Indeed, for fibroblast regulomes, we observe that the set of enhancers positively associated with GHMT is often disjoint from the set of enhancers positively associated with promoter status. Thus, while GHMT induces chromatin opening at regulatory elements across the genome, this opening selectively drives expression of cardiac, rather than fibroblast, genes. PEPR-GNN can use this context-specific responsiveness of gene and enhancer activation to effectively model the behavior of regulomes during reprogramming. Indeed, we find that edges connecting enhancers are among the most informative in our model.

### Limitations of the study

We note several limitations of this study. <u>First</u>, PEPR-GNN narrowly defines regulomes as the perturbations and enhancers that impact a gene’s expression. It excludes other forms of gene regulation including microRNAs, splicing, and post-translational gene regulation. <u>Second</u>, PEPR-GNN discretizes expression phenotypes to simplify our problem to one of classification. However, as gene expression is continuous, the use of arbitrary cutoffs to define binned expression phenotypes can deflate performance measures. <u>Third</u>, PEPR-GNN does not dictate a priori which of the 6 possible edge types between the 4 node types are most informative in the model. While perturbation-enhancer, enhancer-promoter, and promoter-RNA edges represent well-known interactions in enhancer-mediated gene regulation, other edge types can also have biological explanations. Perturbation-promoter edges could represent direct GHMT binding impacting promoter accessibility. Because some portion of the perturbation’s effect on a promoter may occur via enhancer activity, partial confounding of the perturbation-promoter edge type with the corresponding perturbation-enhancer and enhancer-promoter edges, is expected. Perturbation-RNA edges could encapsulate regulation independent of changes in chromatin accessibility. Likewise, enhancer-RNA edges could represent changes in enhancer activity that directly alter gene expression independent of changes in promoter accessibility, such as with promoters with a poised chromatin status^21,22^. Our graphical neural network approach is one of many possible approaches to resolve the relevance of these 6 partially confounding edge types, each with likely gene-dependent impacts.

## Supplemental Figure Legends

**Figure S1, related to Figure 1**

A. Example of enhancer-promoter associations for the Tnnt2 regulome.
B. Perturbation-Enhancer plot analogous to Figure 1G, except colored according to perturbation type to illustrate differing edge distributions. Colors range from NC edges (blue), through intermediate perturbations of 1-3 TFs (green to orange), up to GHMT edges (red).
C. Example regulome architectures for cardiac genes Hcn4 and Ryr2.
D. Example regulome architectures for fibroblast genes Mfap4 and Ccdc80.

**Figure S2, related to Figure 2**

A. Loss per epoch during PEPR-GNN training

**Figure S3, related to Figure 3**

A. Each subplot shows genes colored according to expression response to a specific perturbation combination, in order from NC (0 TFs) up to GHMT (all 4 TFs). B. Genes are colored according to the number of associated enhancers.

## Table Legends

**Table S1**: Sequencing and mapping statistics for the full multiome dataset.

### Key resources table

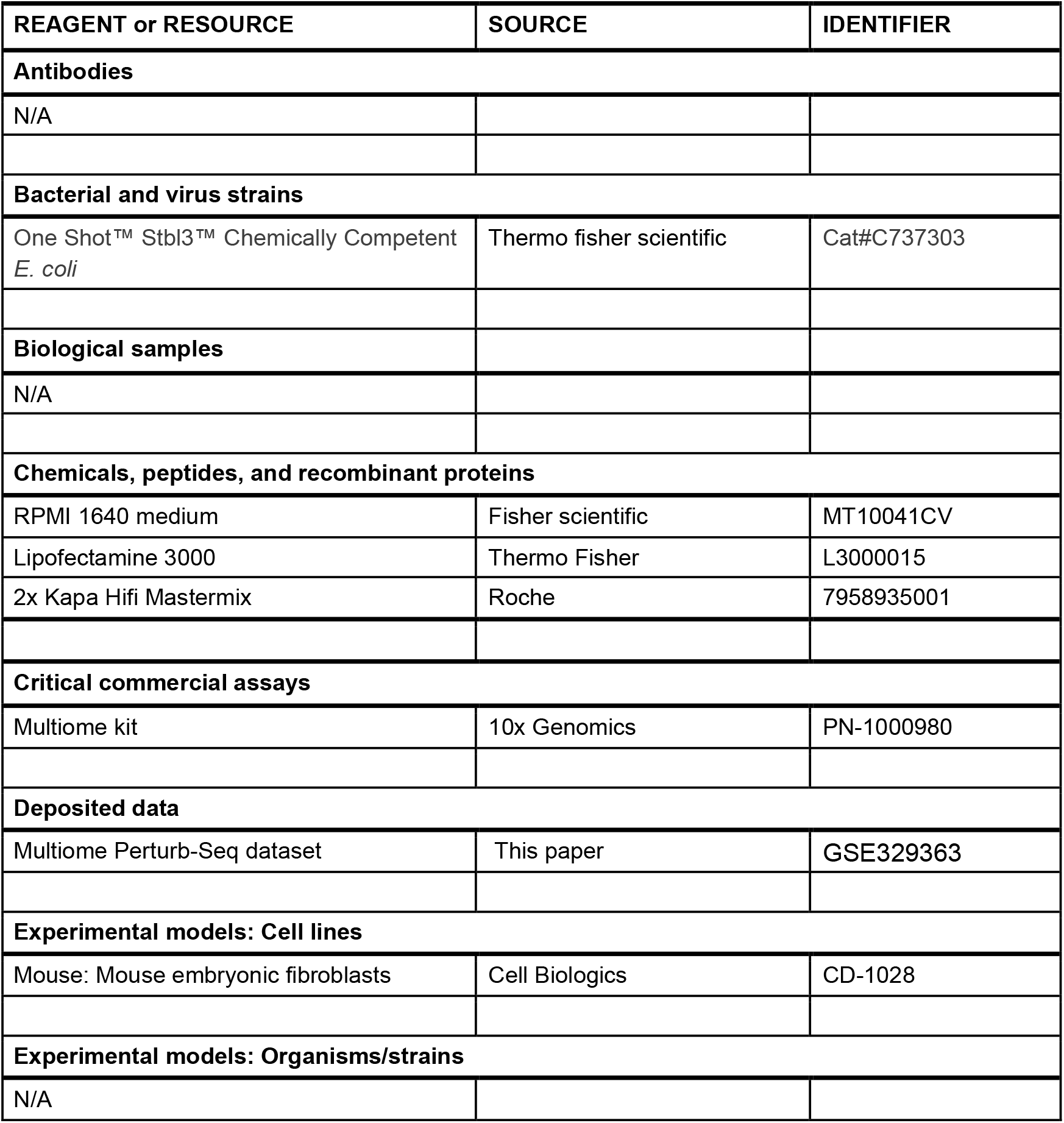
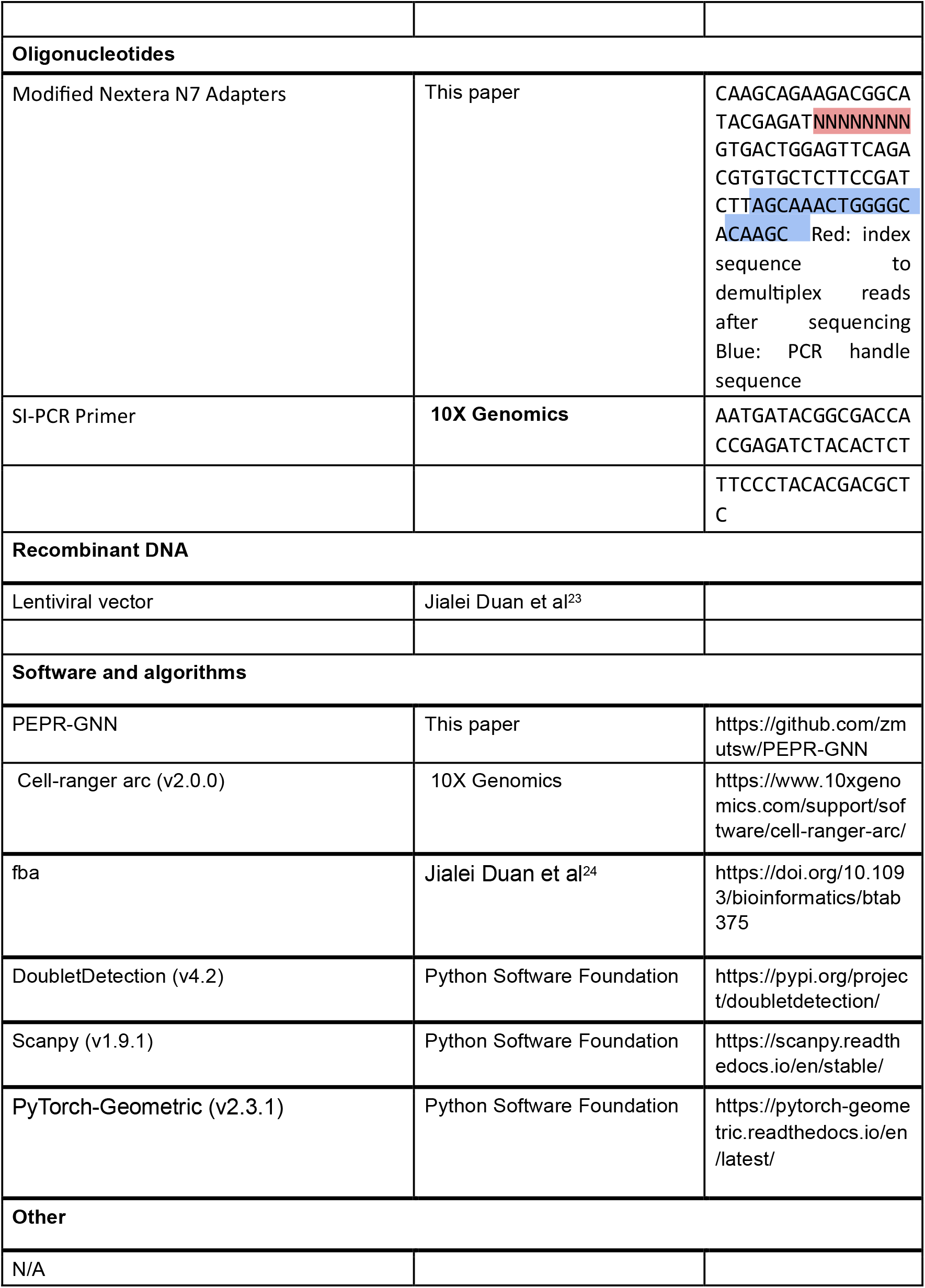

### Resource availability

#### Lead contact

Further information and requests for resources and reagents should be directed to and will be fulfilled by the lead contact, Gary Hon.

#### Data and code availability

The raw and processed data are available at the Gene Expression Omnibus (GEO): GSE329363. This series contains all raw fastq data and processed data including full multiome matrices, perturbation matrices, a table of all significant associations, and a table of properties assigned to each regulome.

Software related to this study are available on GitHub: https://github.com/zmutsw/PEPR-GNN

## STAR Methods

### Cell culture and reprogramming

Commercial Mouse Embryonic Fibroblasts (cMEFs) were grown in growth media (DMEM high-glucose + 10% FBS + 1% penicillin/streptomycin + 1% 100X GlutaMAX). GHMT TFs were synthesized by Twist Bioscience into a lentiviral vector driven by EF1a, followed by lentivirus production with HEK293T cells as described using packaging vectors psPAX2 and pMD2.G^23^. After transduction with lentivirus overnight,the MEFs were incubated in growth media for 14 days. The cells were subsequently washed with DPBS, trypsinized, spun at 500xg for 5 min at 4C, and resuspended in cold growth media.

### Multiome sequencing library preparation

The nuclei were isolated according to 10X protocol CG000366-revB by incubating cells with 0.2X lysis buffer on ice for 15 minutes. The concentration of single nucleus suspension was adjusted to 3200-8000 cells/μL and was loaded on the 10x Genomics Chromium system (10x Genomics, Pleasanton, CA). Single cell RNA-Seq and ATAC-Seq sequencing libraries were constructed following the 10X Chromium Multiome Protocol. To prepare TF perturbation libraries, we used 50-100 ng of purified cDNA amplification product and amplified with primers (SI-PCR Primer from 10X Genomics 3’ single-cell RNA-seq protocol and Modified Nextera N7 Adapter) and 2x Kapa Hifi Mastermix (Roche). We purified the PCR product with a 0.6x SPRIselect beads clean-up. All libraries were sequenced on an Illumina NextSeq sequencer.

### Mapping sequencing data and calling perturbations

Raw sequencing data was mapped using the “count” function from the 10x Genomics package Cell-Ranger arc (v2.0.0) with default parameters. Cell perturbations were called using the kde demultiplex method from fba^24^.

### Filtering and normalization for cells and genes

Raw matrices containing perturbation, ATAC, and RNA, counts per cell were first filtered to remove doublets using the doubletdetection (v4.2) package^25^. Next, cells were removed if they 1) contained more than 25% mitochondrial reads or 2) less than 1,000 total RNA reads, or 3) more than 50,000 total RNA reads. Values in the ATAC matrix were binarized and cells were removed if they differed from the mean number of ATAC reads by more than 2 standard deviations (∼1,000 reads to ∼80,000 reads). Each step removed on the order of 1% of outliers, leaving a total of 8,663 cells detected, with 26,340 genes and 228,876 ATAC peaks.

### Statistical association and graph construction

Annotated transcription start sites that lined up with a called ATAC-Seq peak and RNA were used as the basis for creating a list of 35,070 graphs. All ATAC peaks within 100kb of the TSS included were treated as potential regulatory elements for each graph. This approach gave each graph a node-set of 16 perturbations, a variable set of enhancers, 1 promoter, and 1 RNA. For all edges, fold-change was calculated as the ratio of the mean counts per million (CPM) of the dependent node, conditioned on the presence or absence of the independent node. For edges where the dependent node was RNA, Z-scores were used to account for the large dynamic range. Corresponding p-values were calculated for each edge by 1) creating a 2×2 contingency table based on the presence or absence of the 2 features in question and 2) using the hypergeometric distribution to compare the table’s values against what would be expected if the features were independent.

Node information varied by type. Each perturbation node was encoded as a binary string of the component GHMT factors present in the combination. Enhancer nodes included the distance in bp from the TSS, and both enhancer and promoter nodes included the 6mer count matrix derived from the ATAC peak sequence. In order to reduce redundancy, we counted reverse-compliment motifs together and removed the least variable 50% of motifs, giving a final count of 1122 motifs considered. Finally, graphs with 0 associated enhancers were removed, taking us from 35,070 graphs to 35,038.

### Graph neural network design

We used the pytorch-Geometric (v2.3.1) package for graph data structuring and GNN construction. Our dataset was split into a 4:1 training and testing set, and batched so as to remove the potential for data leakage from overlapping gene loci. Briefly, the PEPR-GNN algorithm consists of 3 modules, each with a GraphConv^26^ layer. The first 2 modules contain ReLu and Dropout functions. During training, the model is fed batched gene graphs as inputs and learns to predict categorized gene expression responses to each perturbation by pooling and transforming outputs before using an Adam optimizer with decreasing learning rate over 2,000 epochs. The loss function is a sigmoid binary cross entropy and is weighted to consider each prediction category (i.e. upregulation, downregulation, or neutral response). Extensive hyperparameter optimization was done, primarily by sequential sweeps while considering various performance metrics such as degree of overfitting and F-score balance across prediction categories.

### Gene embedding analysis

Raw PEPR-GNN outputs (i.e. before pooling and transformation) were used as an information-rich dimensionally uniform representation for each graph. These uniform representations were then fed through a simple dimensionality-reduction pipeline, using the scanpy (v1.9.1) package to perform PCA and tSNE with default paramters. All subsequent embedding analysis was performed with this tSNE embedding, modified as needed to accommodate, e.g., the altered Ttn and Tnnt2 graphs and various visualizations.

### In silico perturbation

For the broad screen a new graph was created for every pair of genes and 6mer motifs (35038*1122) in which the motif counts were set to 0 for all enhancers in the graph. These synthetic graphs were then run through the trained PEPR-GNN model to generate response predictions for each. Prediction scores were then calculated for the synthetic graphs based on the euclidean distance from the synthetic graph to the corresponding real graph. To select candidates for the targeted screen and altered gene embedding, we combed the highest scoring gene-motif pairs for canonical GHMT motifs and cardiac genes. The altered Ttn and Tnnt2 embeddings were generated by setting Tbx5 motif counts to the specified range of values and re-running the gene embedding pipeline on the real graphs in conjunction with the altered graphs.The targeted screen was performed as above except motif values were only incremented by 1 for a single enhancer at a time, giving 1 synthetic graph per enhancer, per candidate gene, from which prediction scores were calculated.

## Acknowledgements

We acknowledge the BioHPC computational infrastructure at UT Southwestern for providing HPC and storage resources that have contributed to the research results reported within this paper. This work is supported by the National Institute of General Medical Sciences (R35GM145235) and the Welch Foundation (I-2103-20250403).

## Author contributions

**Zachary Markham:** Methodology, Software, Formal analysis, Writing - Original Draft, Writing - Review & Editing, Visualization; **Boxun Li**: Investigation; **Landon Nguyen**: Investigation; **Lei Wang:** Investigation; **Nikhil V Munshi:** Conceptualization; **Gary C Hon:** Conceptualization, Writing - Original Draft, Writing - Review & Editing, Supervision, Project administration, Funding acquisition.

## References

1. Ieda, M., Fu, J.-D., Delgado-Olguin, P., Vedantham, V., Hayashi, Y., Bruneau, B.G., and Srivastava, D. (2010). Direct reprogramming of fibroblasts into functional cardiomyocytes by defined factors. Cell 142, 375–386.

2. Takahashi, K., and Yamanaka, S. (2006). Induction of pluripotent stem cells from mouse embryonic and adult fibroblast cultures by defined factors. Cell 126, 663–676.

3. Sekiya, S., and Suzuki, A. (2011). Direct conversion of mouse fibroblasts to hepatocyte-like cells by defined factors. Nature 475, 390–393.

4. Vierbuchen, T., Ostermeier, A., Pang, Z.P., Kokubu, Y., Südhof, T.C., and Wernig, M. (2010). Direct conversion of fibroblasts to functional neurons by defined factors. Nature 463, 1035–1041.

5. Song, K., Nam, Y.-J., Luo, X., Qi, X., Tan, W., Huang, G.N., Acharya, A., Smith, C.L., Tallquist, M.D., Neilson, E.G., et al. (2012). Heart repair by reprogramming non-myocytes with cardiac transcription factors. Nature 485, 599–604.

6. Wada, R., Muraoka, N., Inagawa, K., Yamakawa, H., Miyamoto, K., Sadahiro, T., Umei, T., Kaneda, R., Suzuki, T., Kamiya, K., et al. (2013). Induction of human cardiomyocyte-like cells from fibroblasts by defined factors. Proc Natl Acad Sci U S A 110, 12667–12672.

7. Wang, H., Yang, Y., Liu, J., and Qian, L. (2021). Direct cell reprogramming: approaches, mechanisms and progress. Nat Rev Mol Cell Biol 22, 410–424.

8. Zhao, H., Zhang, Y., Xu, X., Sun, Q., Yang, C., Wang, H., Yang, J., Yang, Y., Yang, X., Liu, Y., et al. (2021). Sall4 and Myocd Empower Direct Cardiac Reprogramming From Adult Cardiac Fibroblasts After Injury. Front Cell Dev Biol 9, 608367.

9. Zhang, Z., Zhang, W., and Nam, Y.-J. (2019). Stoichiometric optimization of Gata4, Hand2, Mef2c, and Tbx5 expression for contractile cardiomyocyte reprogramming. Scientific Reports 9, 14970.

10. Hashimoto, H., Wang, Z., Garry, G.A., Malladi, V.S., Botten, G.A., Ye, W., Zhou, H., Osterwalder, M., Dickel, D.E., Visel, A., et al. (2019). Cardiac Reprogramming Factors Synergistically Activate Genome-wide Cardiogenic Stage-Specific Enhancers. Cell Stem Cell 25, 69–86.e5.

11. Kipf, T.N., and Welling, M. (2016). Semi-Supervised Classification with Graph Convolutional Networks.

12. Stewart, A.J., Hannenhalli, S., and Plotkin, J.B. (2012). Why transcription factor binding sites are ten nucleotides long. Genetics 192, 973–985.

13. GitHub - pyg-team/pytorch_geometric: Graph Neural Network Library for PyTorch GitHub. https://github.com/pyg-team/pytorch_geometric.

14. Hinton, G.E., Srivastava, N., Krizhevsky, A., Sutskever, I., and Salakhutdinov, R.R. (2012). Improving neural networks by preventing co-adaptation of feature detectors.

15. Website 10.1016/j.neucom.2022.06.111.

16. Terven, J., Cordova-Esparza, D.M., Ramirez-Pedraza, A., Chavez-Urbiola, E.A., and Romero-Gonzalez, J.A. (2023). Loss Functions and Metrics in Deep Learning. 10.1007/s10462-025-11198-7.

17. Riching, A.S., Zhao, Y., Cao, Y., Londono, P., Xu, H., and Song, K. (2018). Suppression of Pro-fibrotic Signaling Potentiates Factor-mediated Reprogramming of Mouse Embryonic Fibroblasts into Induced Cardiomyocytes. J Vis Exp. 10.3791/57687.

18. Zhao, Y., Londono, P., Cao, Y., Sharpe, E.J., Proenza, C., O’Rourke, R., Jones, K.L., Jeong, M.Y., Walker, L.A., Buttrick, P.M., et al. (2015). High-efficiency reprogramming of fibroblasts into cardiomyocytes requires suppression of pro-fibrotic signalling. Nat Commun 6, 8243.

19. Jolma, A., Yan, J., Whitington, T., Toivonen, J., Nitta, K.R., Rastas, P., Morgunova, E., Enge, M., Taipale, M., Wei, G., et al. (2013). DNA-binding specificities of human transcription factors. Cell 152, 327–339.

20. Website JASPAR 2026: expansion of transcription factor binding profiles and integration of deep learning models, Nucleic Acids Research, Volume 54, Issue D1, 6 January 2026, Pages D184–D193, 10.1093/nar/gkaf1209.

21. Rada-Iglesias, A., Bajpai, R., Swigut, T., Brugmann, S.A., Flynn, R.A., and Wysocka, J. (2011). A unique chromatin signature uncovers early developmental enhancers in humans. Nature 470, 279–283.

22. Pachano, T., Sánchez-Gaya, V., Ealo, T., Mariner-Faulí, M., Bleckwehl, T., Asenjo, H.G., Respuela, P., Cruz-Molina, S., Muñoz-San Martín, M., Haro, E., et al. (2021). Orphan CpG islands amplify poised enhancer regulatory activity and determine target gene responsiveness. Nature Genetics 53, 1036–1049.

23. Duan, J., Li, B., Bhakta, M., Xie, S., Zhou, P., Munshi, N.V., and Hon, G.C. (2019). Rational Reprogramming of Cellular States by Combinatorial Perturbation. Cell Rep. 27, 3486–3499.e6.

24. Website Duan, J., & Hon, G. (2021). FBA: feature barcoding analysis for single cell RNA-Seq. Bioinformatics, 37(22), 4266–4268. 10.1093/bioinformatics/btab375.

25. Website Adam Gayoso, & Jonathan Shor. (2025). JonathanShor/DoubletDetection: doubletdetection v4.3.0.post1 (v4.3.0.post1). Zenodo. 10.5281/zenodo.14827937.

26. Morris, C., Ritzert, M., Fey, M., Hamilton, W.L., Lenssen, J.E., Rattan, G., and Grohe, M. (2018). Weisfeiler and Leman Go Neural: Higher-order Graph Neural Networks.

