## Supplementary figures and images for "PEPR-GNN: Perturbation-Enhancer-Promoter-RNA Graph Neural Networks for Multiome Perturb-Seq modeling of regulomes"

### Figure S1

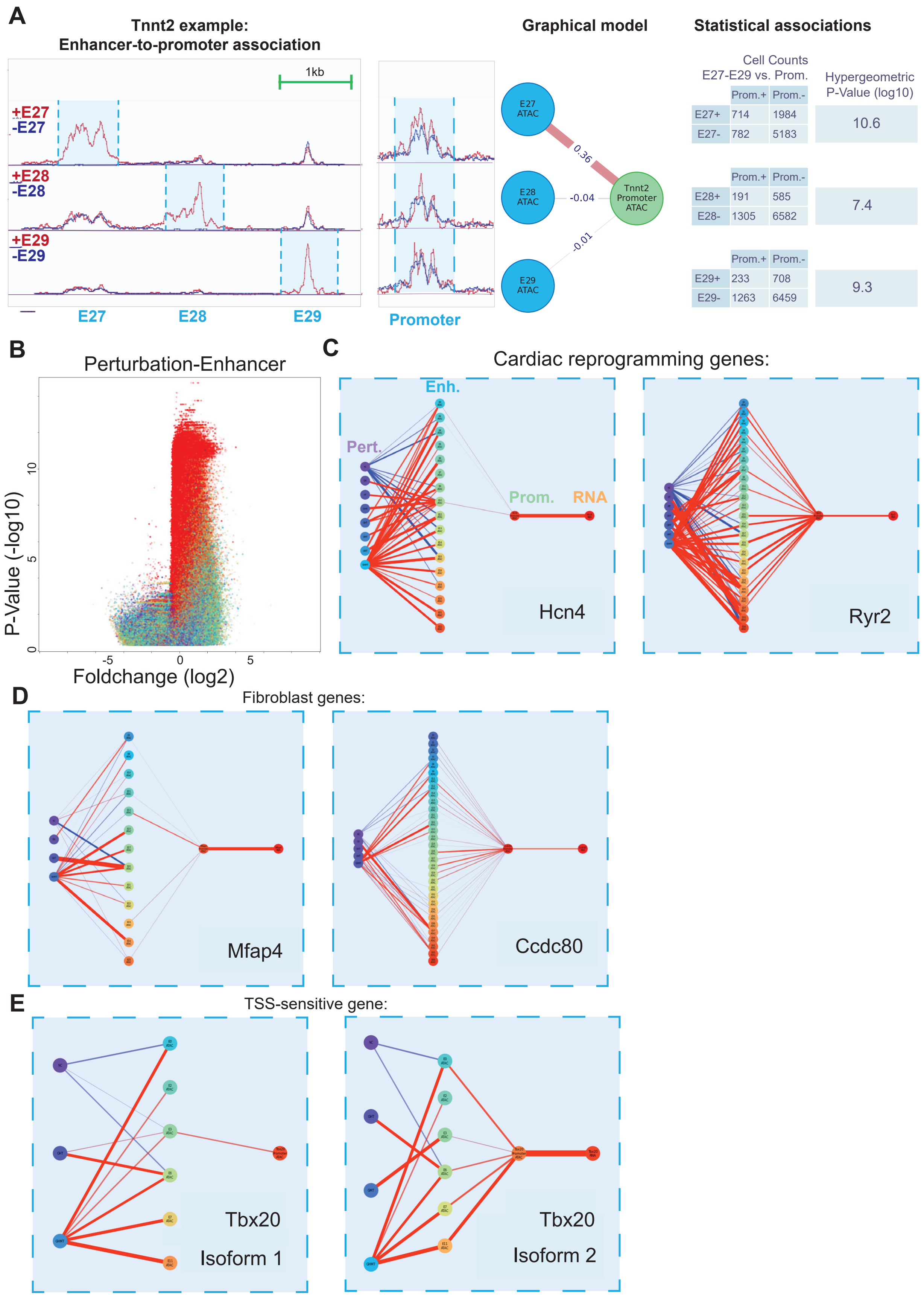

### Figure S2

A

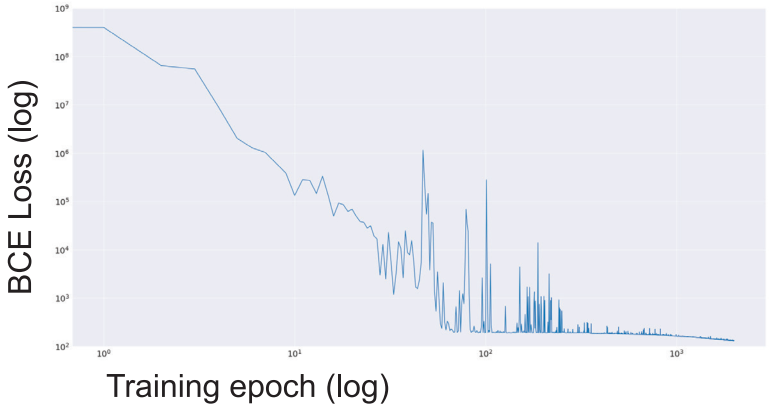

### Figure S3

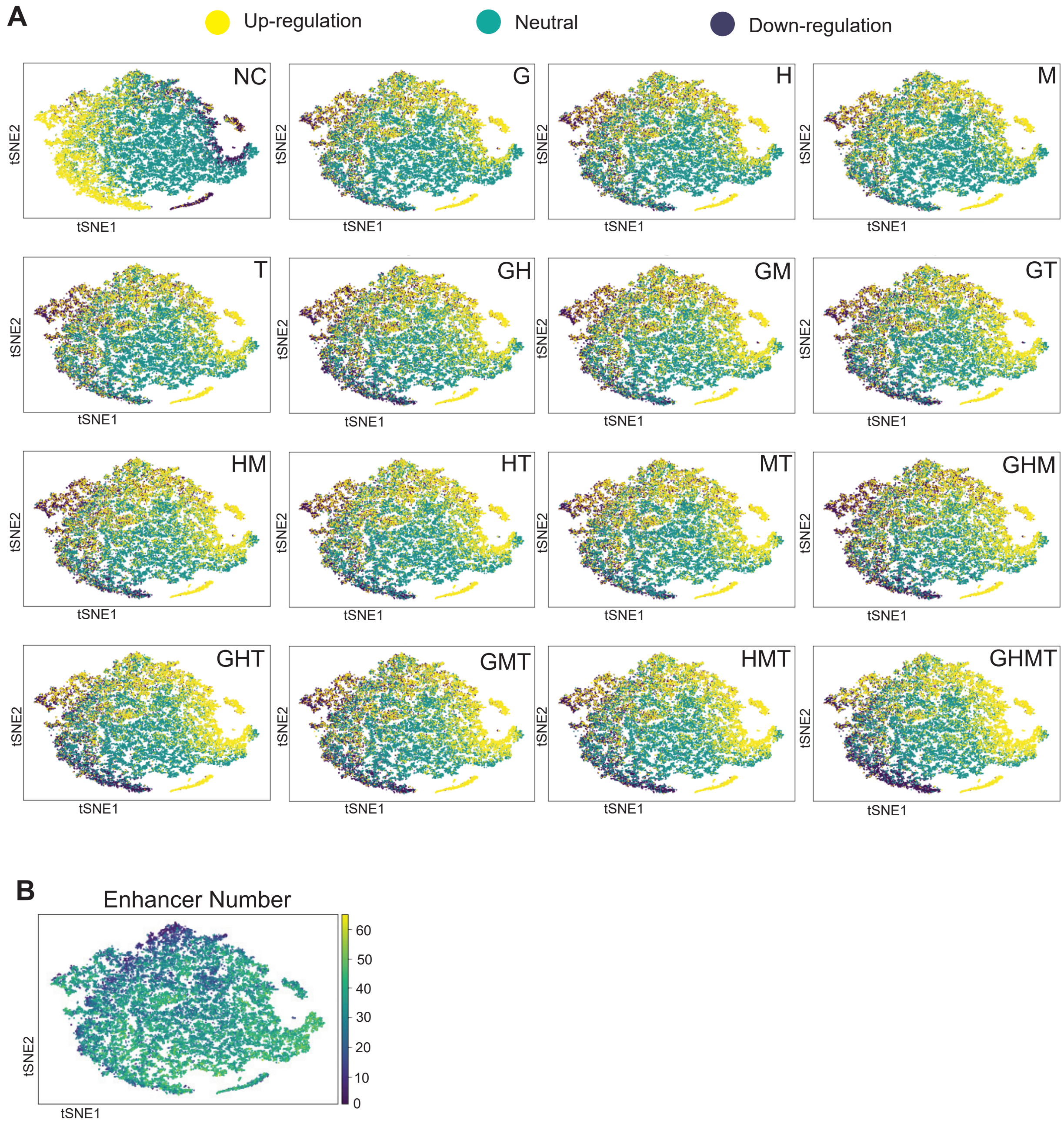
